# Phase-based cortical synchrony is affected by prematurity

**DOI:** 10.1101/2021.02.15.431226

**Authors:** Pauliina Yrjölä, Susanna Stjerna, J. Matias Palva, Sampsa Vanhatalo, Anton Tokariev

## Abstract

Inter-areal synchronization by phase-phase correlations (PPC) of cortical oscillations mediates many higher neurocognitive functions, which are often affected by prematurity, a globally prominent neurodevelopmental risk factor. Here, we used electroencephalography (EEG) to examine brain-wide cortical PPC networks at term-equivalent age, comparing human infants after early prematurity to a cohort of healthy controls. We found that prematurity affected these networks in a sleep state-specific manner, and the differences between groups were also frequency-selective, involving brain-wide connections. The strength of synchronization in these networks was predictive of clinical outcomes in the preterm infants. These findings show that prematurity affects PPC networks in a clinically significant manner suggesting early functional biomarkers of later neurodevelopmental compromise to be used in clinical and translational studies after early neonatal adversity.

## Introduction

Approximately 10% of infants are born preterm, which inflicts lifelong disabilities in many key brain functions, including vision, learning, and language processing (Johnson & Marlow, 2017; WHO, 2012). Many of these functional abnormalities arise from the impacts that prematurity has on neuronal networks. Recent studies have demonstrated both structural (Batalle et al., 2017; Guo et al., 2017) and functional (Tokariev et al., 2019a; Tokariev et al., 2019b; Tóth et al., 2017) effects of prematurity, some of which are shown to predict later neurodevelopmental outcomes.

Prematurity implies that the infants spend a part or all of their third trimester of gestation in an unnatural environment, *ex utero.* This time window is known to be characterized by the growth of brain networks driven by a combination of genetic and activity-dependent mechanisms (Luhmann et al., 2016; Molnár et al., 2020). The early cortical activity can be recorded with scalp electroencephalography (EEG) and consists of spontaneous intermittent bursts, which provide an early mechanism for inter-areal temporal correlations and define functional cortical networks (Vanhatalo & Kaila, 2006). Therefore, the early cortical activity is a driver, guide, and biomarker of the development of brain networks.

The functional cortical networks can be characterized by quantifying relationships between phase or amplitude attributes of neural signals from distinct brain regions. Prior research on neonatal EEG (Omidvarnia et al., 2014; Tokariev et al., 2019a; Tokariev et al., 2019b) have often focused on the amplitude–amplitude correlations (AACs) that reflect co-modulation of overall neuronal activity and gross cortical excitability over periods of seconds (Engel et al., 2013; Hipp et al., 2012; Palva & Palva, 2011; Tewarie et al., 2019). The other commonly used measure of neuronal interactions is phase–phase correlation (PPC) that is considered to reflect a spatiotemporally accurate mechanism of inter-areal communication. PPC is thought to arise from subsecond timing relationships in neuronal spiking (Palva & Palva, 2011; Vidaurre et al., 2018; Womelsdorf et al., 2007), hence being able to support dynamic integration in neuronal ensembles underlying several higher-level brain functions (Bressler & Menon, 2010; Palva & Palva, 2011). Moreover, it is now well known that the brain operates concurrently at multiple frequencies, giving rise to multiplex networks shaped by concerted actions of different coupling mechanisms in several frequency bands (De Domenico et al., 2016; Siebenhühner et al., 2016).

Our recent study suggests that PPC networks link to neurological performance (Tokariev et al., 2019b). However, the spatial and spectral extent of these findings, as well as their clinical correlates have remained unclear. Here, we aimed to assess how the large-scale cortical PPC networks are affected by preterm birth of human infants. We analysed EEG recordings from a large cohort of preterm and healthy control infants using an infant-specific source modelling-based analysis pipeline that allows non-invasive assessment of functional networks at the level of cortical sources. We asked whether prematurity leads to changes in the cortical networks that are linked to sleep state, brain area or oscillation frequency. Moreover, we wanted to study if prematurity-related changes in cortical networks would have clinical significance, i.e., be predictive of clinical neurological performance of the infants by the time of recording and/or later during childhood.

## Results

To characterize effects of prematurity on early cortical networks, we recorded multichannel scalp EEG at term-equivalent age from a group of infants born extremely preterm (EP, N = 46), as well as from a group of full-term healthy controls (HC, N = 67). The PPC cortical networks were computed from source-reconstructed EEG data for active (AS) and quiet sleep (QS) within 21 narrow frequency bands covering the physiologically relevant range of 0.4–22 Hz (Vanhatalo et al., 2005). To evaluate the impact of prematurity on cortical networks as a function of frequency, we estimated the extent of significant network differences between EP and HC groups at each specific frequency. Finally, to define patterns which are linked to infant neurological performance, we correlated the PPC networks of the EP group to key neurological scores at term age and neurocognitive assessments at two years of age. We described the extent of patterns that are different between groups or correlate to outcomes as a fraction of statistically significant connections *(K)* relative to the whole network (Palva et al., 2010). The overall analytical flow is shown on Figure 1 and described in detail in the *Methods* section.

**Figure 1.**
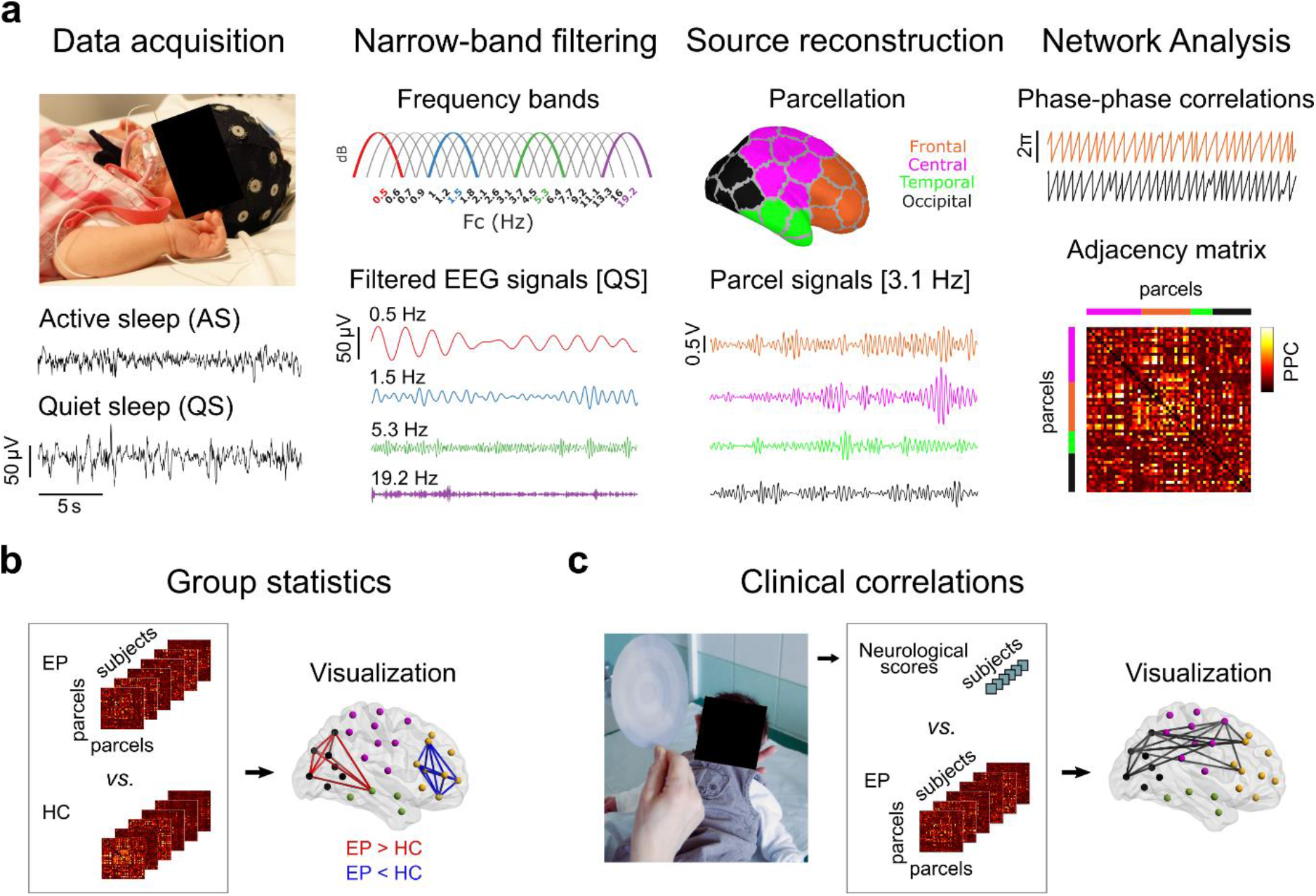
Outline of the study design and analyses. **(a)** EEG recordings of day-time sleep were acquired from early preterm (EP) and healthy control (HC) cohorts. The recordings were classified into active (AS) and quiet sleep (QS), and 5-minute-long epochs were constructed for both sleep states. The selected epochs were filtered into 21 narrow frequency bands of semi-equal length on a logarithmic scale and converted to cortical source signals applying a realistic infant head model with 58 cortical parcels. Functional connectivity analysis was applied on the parcel signals by computing phase-phase correlations (PPCs) with the debiased phase lag index, yielding subject-specific connectivity matrices for both sleep states and all frequency bands. **(b)** Statistical group differences in connectivity strength were computed (Wilcoxon rank sum test) for both sleep states and each frequency band. The edges portraying significant differences for two contrasts EP > HC (red) and EP < HC (blue) were then visualized. **(c)** Finally, correlations of PPC strengths to newborn neurological and 2-year neurocognitive assessment scores were investigated (Spearman correlation). The edges related to significant clinical correlation were visualized.

### PPC networks are affected by prematurity in a frequency-specific manner

We found broad sleep- and frequency- specific differences in PPC networks between EP and HC infants (Figure 2). During AS, the most extensive group differences were observed within the theta frequency band (peak at Fc = 5.3 Hz; Figure 2a) with stronger connections in the EP group *(K* = 10%, *p* < 0.01, *q* = 0.01) that were uniformly distributed over the whole cortex (Figure 2b) and preferentially long-range. Smaller subnetworks (*K* = 2–4%, *p* < 0.01, *q* = 0.01) of both increased and decreased connectivity in EP infants were at delta frequencies (1.8–3.1 Hz) covering central and temporal regions. During QS, the most prominent group differences were within the delta band: The EP infants exhibited stronger connectivity (*K* = 10% at 1.8 Hz, *p* < 0.01, *q* = 0.01) in mostly long-range connections between frontal and occipital lobes, while there were weaker short-range connections (*K* = 8% at 1.2 Hz, *p* < 0.01, *q* = 0.01) within the frontal lobe and a few projections to the parietal lobe. Networks at alpha and beta frequencies were suppressed in the EP infants during both sleep states, and they mainly involved basal connections linking occipital cortices to frontal and temporal areas.

**Figure 2.**
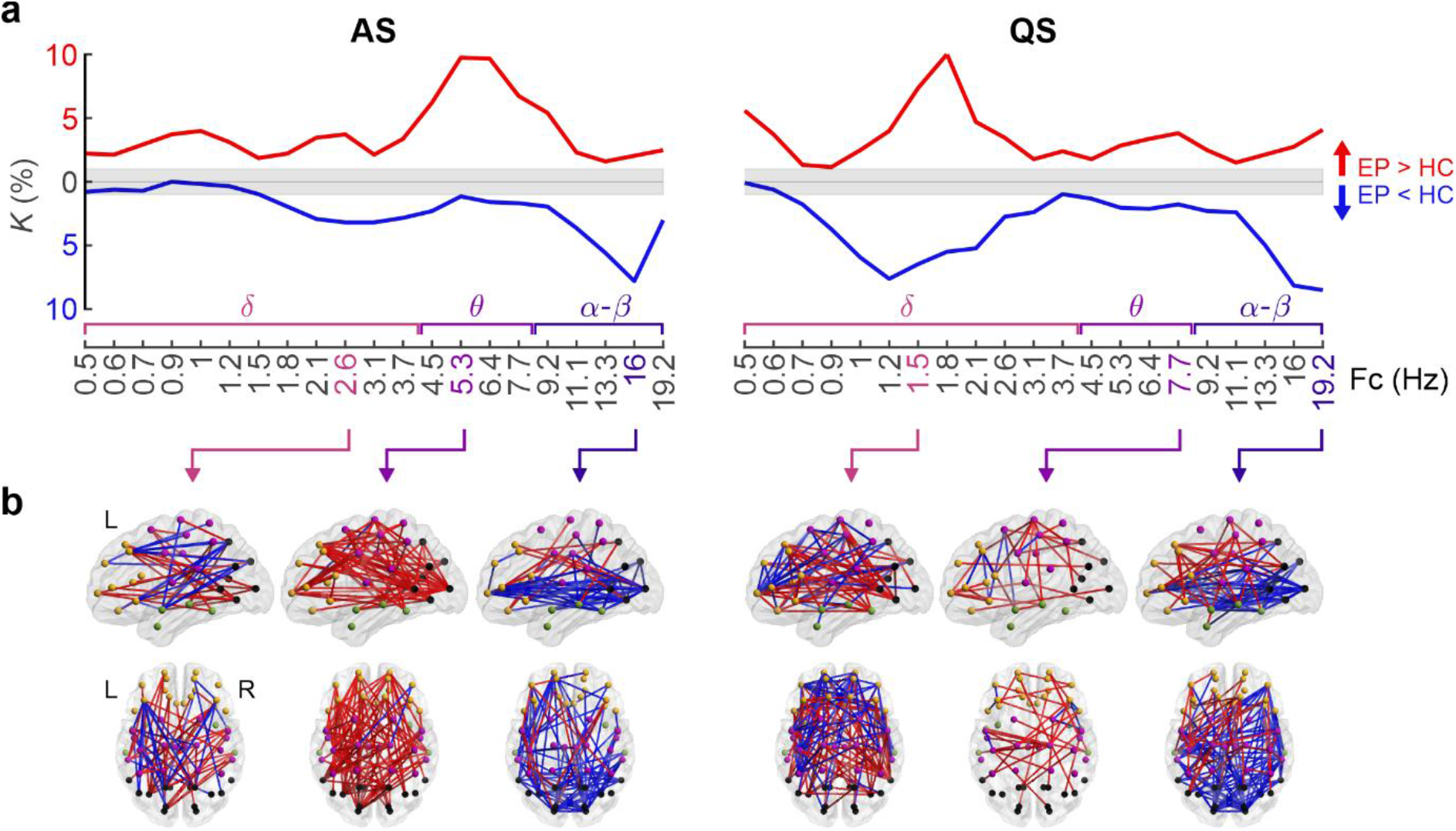
Effects of prematurity on cortical PPC networks. **(a)** Network density *(K)* of significant PPC group differences (two one-tailed Wilcoxon rank-sum tests, α = 0.01) during active sleep (AS, left) and quiet sleep (QS, right) as a function of frequency. Networks that are stronger in EP (EP > HC) are shown in red, whereas networks with suppressed connectivity in EP (EP < HC) are presented in blue. The grey shaded area depicts the boundaries of the *q*-level showing the potential level of false discoveries (*q* = 0.01). The data presented in the figure is provided in Figure 2—source data 1 and matrices of the *p*-values and effect sizes of all networks in Figure 2—source data 2. **(b)** Spatial visualizations present PPC network comparisons at the frequencies with the most extensive group differences. The color coding of the networks (red, blue) is equal to that of (a).

No significant correlations (Spearman, two-tailed test, α = 0.05, Benjamini-Hochberg correction) were found between mean connectivity strength and age per frequency. Effect sizes, computed by the rank-biserial correlation over each significant network, are presented in Figure 2—figure supplement 1. The spatial differences in PPC networks between groups for all frequency bands are shown at Figure 2—figure supplement 2. We also validated the results with an alternative analysis using network-based statistics (NBS), (Zalesky et al., 2010), with two one-tailed tests (for details, see *Methods*), and we found strikingly similar spectral and spatial patterns in group comparisons (Figure 2—figure supplement 3). The findings together suggest that exposure to prematurity affects the organization of cortical PPC networks at limited oscillatory frequencies.

### Connectivity strength correlates with neurological performance in preterm infants

Next, we studied how the strength of cortical PPC networks correlates to neurological performance at the time of newborn EEG recordings. To this end, we correlated the connectivity strengths of each PPC network connection (N = 1128) of the EP group to the neurological performance of the corresponding infants, assessed using compound scores C1 and C2, which are associated with later motor and cognitive outcomes, respectively (Tokariev et al., 2019b). We estimated the fraction of significantly correlated connections using a density measure (*K*) across the whole frequency domain, and visualized the networks showing broad spatial effects (Figure 3).

**Figure 3.**
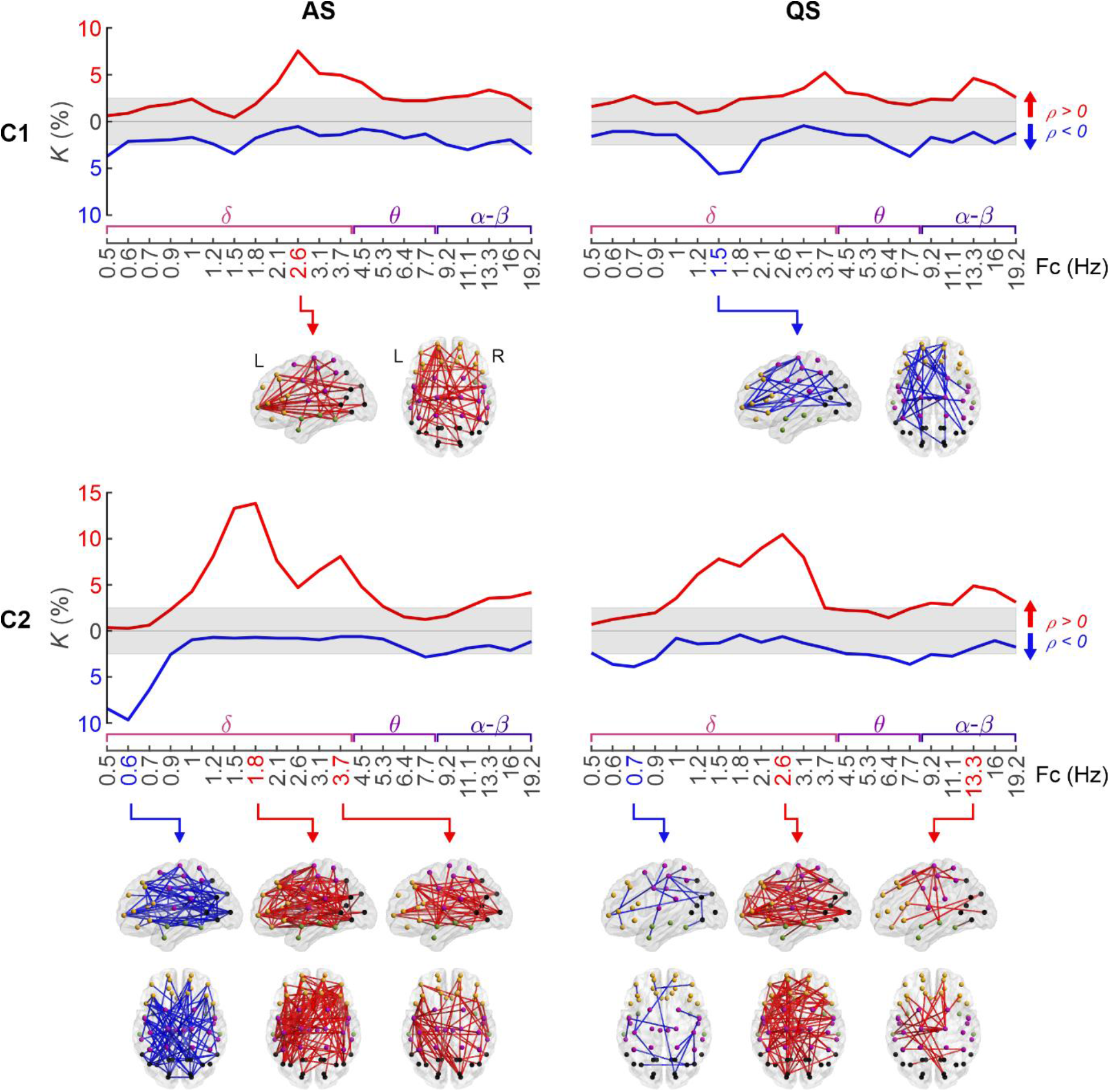
PPC networks of ex-preterm infants at term age predict neurological outcome. Density (*K*) of PPC patterns that associate to neurological scores C1 and C2 (Spearman, twotailed test with conceptional age as a covariate, α = 0.05) as a function of frequency. The grey shaded area depicts the FDR boundaries (*q* = 0.05). The opaque brains show the spatial distributions of networks taken at the most characteristic peaks of the density curves. Red coloring pictures networks with positive correlation *(ρ* ≥ 0), while blue coloring shows negatively correlated connections (*ρ* < 0) in both the graphs and 3-dimensional plots. The graph data is provided in Figure 3—source data 1 and the full *p*-value and effect size matrices in Figure 3—source data 2.

The C2 score was positively correlated with an extensive pattern at higher delta frequencies in both sleep states (AS: 1.2–4.5 Hz, *K* = 5–14%, QS: 1.2–3.1 Hz, *K* = 6–10%, *p* < 0.05, *q* = 0.05). The corresponding spatial patterns incorporated broad networks linking multiple distal areas. In contrast, the C1 score showed only mildly elevated density, or small networks, with positive correlation at 2.6–4.5 Hz during AS (*K* = 4–8%, *p* < 0.05, *q* = 0.05), and a negative correlation at 1.5–1.8 Hz during QS (*K* = 5–6%, *ρ* < 0.05, *q* = 0.05). Effect sizes (mean of Spearman *ρ* over the positive and negative correlation networks) are depicted in Figure 3— figure supplement 1. A comparable analysis for healthy controls (Figure 3—figure supplement 2) showed only a few negative correlations between edge strength and neurological scores. The spatial distributions for all investigated frequency bands are presented in Figure 3—figure supplement 3 for C1 and Figure 3—figure supplement 4 for C2.

These findings together suggest that the relationship between cortical networks and neurological performance is affected by prematurity. The EP infants exhibit brain-wide relationships between cortical networks and neurological performance, which is not seen in the HC infants.

### Correlation of functional connectivity and neurological performance extends to longterm neurocognitive outcomes

Finally, we examined the relation of PPC networks to long-term neurocognitive development, assessed at 2 years of age using standardized Bayley (Bayley, 2006) and Griffiths (Huntley, 1996) scores. Akin to our analysis above on newborn clinical performance, we also correlated the strength of individual connections in the PPC networks of the EP infants to their clinical outcome measures at two years of age. Most of the significant correlations emerged for visual, motor, cognitive, and language comprehension scores at lower frequencies (Figure 4), forming mostly spatially constrained patterns.

**Figure 4.**
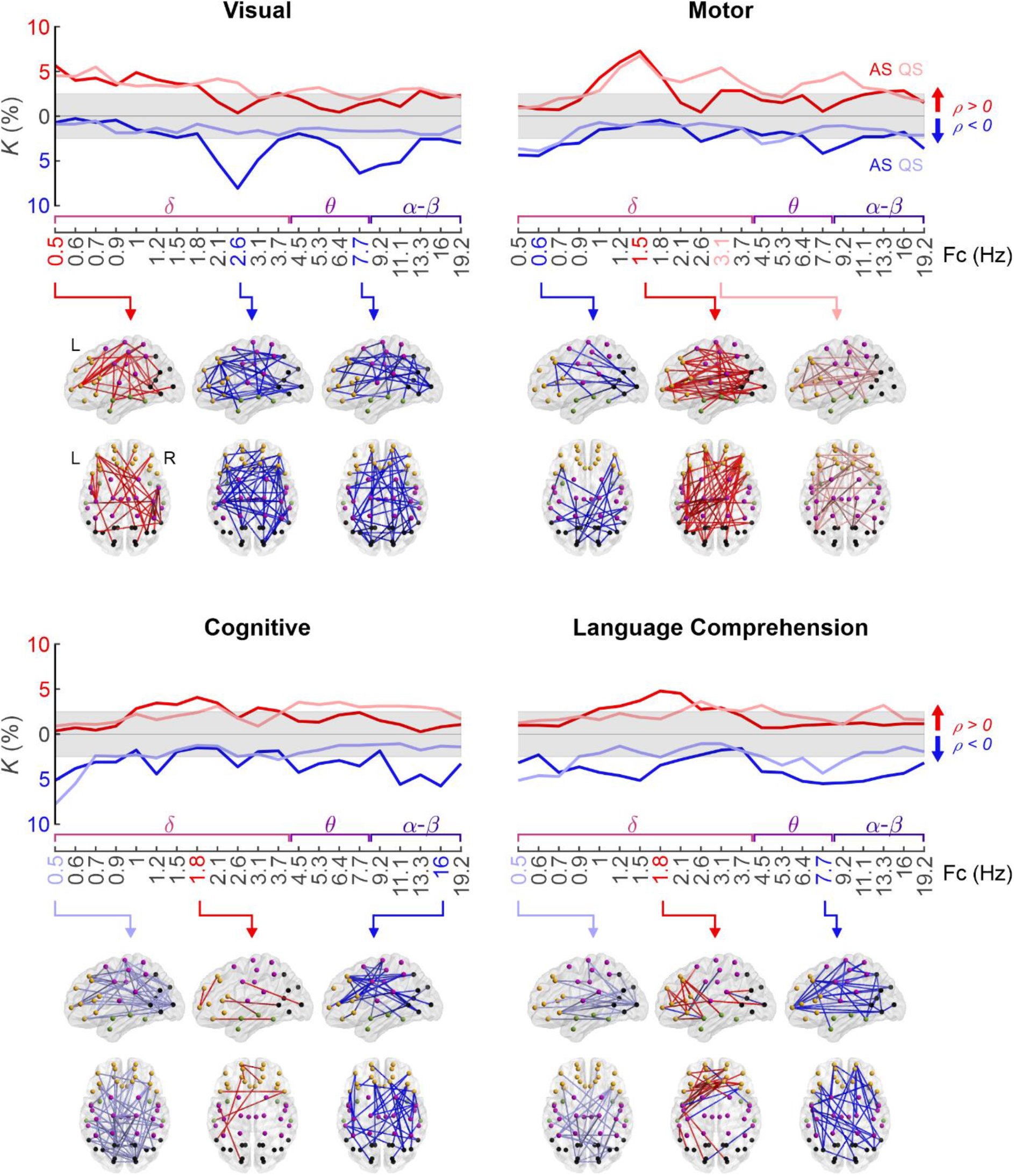
Correlation of PPC network strength to 2-year neurocognition. The upper graphs show the frequency-wise summary of the proportion of network edges (*K*) that show a significant correlation between PPC strength and the given neurocognitive performance score (Spearman, two-tailed test with conceptional age as a covariate, α = 0.05). The FDR (*q* = 0.05) boundaries are depicted as a grey shaded area. The strongest peaks in these plots were selected for the 3-dimensional visualisations of networks as indicated with arrows. Colour coding represents the sign of correlation (red: *ρ* ≥ 0, blue *ρ* < 0) and hues represent sleep states (dark: AS, light: QS) in the graphs and the spatial visualisations. The data displayed in the curves is provided in Figure 4—source data 1 and the *p*-value and effect size matrices from which the graphs were created in Figure 4—source data 2.

Visual scores correlated positively with PPC at Fc = 0.5 Hz during both sleep states (*K* = 5–6%, *p* < 0.05, *q* = 0.05), involving networks from left frontal and right occipital regions. There were also some negative correlations between visual scores and PPC during AS (*K* = 6–8%, *p* < 0.05, *q* = 0.05), located mostly in the frontal regions (Fc = 2.6 Hz) or occipital regions (Fc = 7.7 Hz).

Motor scores featured a prominent positive correlation during both sleep states at Fc = 1.5 Hz (*K* = 7%, *p* < 0.05, *q* = 0.05) with a broad spatial distribution over several cortical regions. We also found a somewhat smaller extent network with a positive correlation to motor score during QS at a slightly higher frequency (Fc = 3.1 Hz; *K* = 5%, *p* < 0.05, *q* = 0.05). Finally, a subset of occipital interhemispheric connections showed negative correlation to motor scores during both sleep states at the lowest frequencies (Fc = 0.6 Hz; *K* = 4%, *p* < 0.05, *q* = 0.05).

The cognitive performance score showed negative correlations at low (AS and QS; Fc = 0.5 Hz; *K* = 5–8%, *p* < 0.05, *q* = 0.05) and high frequencies (AS only; Fc = 16 Hz; *K* = 6%, *p* < 0.05, *q* = 0.05), involving networks that connect frontal, parietal, and occipital regions. A smaller network displayed positive correlations with cognitive performance during AS at Fc = 1.8 Hz (*K* = 4%, *p* < 0.05, *q* = 0.05), involving mostly frontal connections.

Language comprehension was strongly and positively correlated to PPC strength in AS (peak at Fc = 1.8 Hz, *K* = 5%, *p* < 0.05, *q* = 0.05), involving networks from the left temporal to frontal regions, aligning well with the cortical areas that are known to participate in language comprehension (Tremblay & Dick, 2016). A negative correlation between PPC strength and language comprehension scores was present during QS in the basal long-range connections at lower frequencies (Fc = 0.5 Hz, *K* = 5%, *p* < 0.05, *q* = 0.05), as well as diffuse brain-wide network at mid-frequencies during AS (Fc = 7.7 Hz, *K* = 4%, *p* < 0.05, *q* = 0.05).

Effect sizes were computed as the mean of Spearman *ρ* of the positive and negative networks separately and are presented as a function of frequency in Figure 4—figure supplement 1. The spatial distributions at all investigated frequency ranges are shown in Figure 4—figure supplement 2–5.

## Discussion

Our study shows that spontaneous cortical activity in the human infants exhibits large-scale PPC structures, that are spectrally and spatially selective and co-vary with vigilance states. Moreover, we show that the globally most significant clinical risk factor, preterm birth (WHO, 2012), leads to frequency-selective changes in these networks, which correlate to neurocognitive performance of the affected individuals. Our work employed novel realistic cortical source reconstruction and independent parallel analyses to validate the results on clinical network correlations. Our findings are broadly consistent with recent work on adults showing that multiple frequency-specific PPC networks coexist (De Domenico et al., 2016; Siebenhühner et al., 2016; Vidaurre et al., 2018; Yu et al., 2017) and correlate with normal and pathological behaviour (Siebenhühner et al., 2016; Yu et al., 2017). Our work extends prior studies reporting prematurity effects on the temporally loose amplitude correlations (Omidvarnia et al., 2014; Tokariev et al., 2019a); here we provide evidence that the cortico-cortical interactions in newborn infants are already accurate enough to give rise to spectrally and spatially specific PPC structures, and pathological effects therein.

It has recently become clear that brain function relies on several co-existing frequency-specific PPC networks, which are reported to show temporal dynamics between awake states in the adults (Siebenhühner et al., 2016; Vidaurre et al., 2018) or between sleep states in the neonatal studies (Tokariev et al., 2019b; Tokariev et al., 2016b). Here, we show that medical adversities can affect these PPC networks in a selective manner, at preferential frequencies, and with preferential spatial distributions, as well as differing between vigilance states. For instance, prematurity caused an increase in middle frequency PPC in long-range connections throughout the brain, while the changes in higher frequencies were more localized in the middle and long-range connections in the basal brain areas. These effects were more pronounced during active sleep for the middle frequencies, while high frequency findings were essentially similar between sleep states. The findings are compatible with a notion that the functional significance of frequency-specific PPC networks depends on their context, the brain state, in addition to their given frequency.

Our present findings extend the long-held clinical tradition where quiet sleep is considered to be the most sensitive state in disclosing effects of prematurity in the EEG records. In the clinical visual review, the EEG signal is considered to exhibit dysmature/immature features (Lombroso, 1979; Tharp, 1990), and the most robust feature is augmented “interhemispheric asynchrony”, or temporal non-overlap between cortical bursting (Koolen et al., 2014; Räsänen et al., 2013). While the clinically perceived interhemispheric asynchrony considers quiet sleep and amplitude correlations only, here we show that robust prematurity effects are also seen in the PPC networks, and they are clear during active sleep. Moreover, the functional significance of the PPC network during active sleep is shown by their pronounced correlations to subject-level clinical performance.

The strength of PPC connectivity in several brain-wide subnetworks was found to correlate to infants’ neurological performance at term age, which extends prior reports on clinical correlations to frontally connected delta frequency networks (Tokariev et al., 2019b). Clinical correlations were clearly widest for the C2 composite score which emphasizes features of newborn performance that pre-empt later cognitive development (Tokariev et al., 2019b). Comparison to neurocognitive performance at 2 years of age showed also several albeit smaller PPC subnetworks with significant correlations. PPC connectivity relies on a temporally accurate neural communication that requires sufficiently matured cortico-cortical pathways (Palva & Palva, 2011; Womelsdorf et al., 2007). The previously described diffuse and extensive white matter abnormalities after prematurity (Dimitrova et al., 2020) may provide a straightforward histological underpinning for the changes, especially the observed decrease in higher frequency PPC networks.

While our findings suggest clinically meaningful functions for the herein characterized PPC networks in the EP infants, it was somewhat unexpected that comparable correlations were not found in the group of healthy control infants. That observation suggests an altered relationship between PPC networks and neurocognitive phenotypes, which calls for a reasonable mechanistic explanation. It is possible that the network-phenotype relationship becomes amplified in the preterm cohort that is known to exhibit considerable variation in their histological maturation (Dimitrova et al., 2020). Prior studies have shown brain-wide effects of prematurity on the histological structures of white matter tracts (Dimitrova et al., 2020) and these changes were shown to correlate with several characteristics of newborn or later neurocognitive performance (Girault et al., 2019; Stjerna et al., 2015; Toulmin et al., 2020; Vollmer et al., 2017). An alternative mechanism is that the effects found in EP infants reflect a transient network immaturity (Lombroso, 1979; Tharp, 1990) that would catch up during later development. Testing this hypothesis would need repeated EEG network studies in the EP infants near term-equivalent age to show a developmental catch up in the PPC networks (Tokariev et al., 2016b) or in other functional brain age (Stevenson et al., 2020).

The present results suggest a clinically meaningful effect on PPC networks that could potentially serve as a functional biomarker to, *e.g.,* benchmark early therapeutic interventions (Ewen et al., 2019; Sahin et al., 2020). Our current study needs to be considered as observational work that identified putative analysis pipelines and network markers. Future prospective studies on larger cohorts are needed for their validation, and to define the perceived added value of network assessment from the perspective of monitoring early neurodevelopment and benchmarking early therapeutic interventions. In addition, the hereby demonstrated network effects may offer a unique translational bridge: The PPC networks could be used as a functional benchmark for establishing the clinical neurodevelopmental relevance of preclinical models of human prematurity.

## Methods

The full study pipeline can be viewed in Figure 1.

### Subjects

The dataset included N = 46 early preterm (EP) and N = 67 healthy control (HC) infants. The conceptional ages (CA) at birth of the EP group (mean ± standard deviation, SD) were 24.4 ± 1.2 weeks and those of the HC group were 38.4 ± 1.1 weeks. This dataset was collated from cohorts that have been published in previous studies (Omidvarnia et al., 2014; Tokariev et al., 2016b; Tokariev et al., 2019a; Tokariev et al., 2019b). The study design was approved by the Ethics Committee of the Helsinki University Central Hospital and informed consent was obtained from a parent or guardian for each subject.

### EEG recordings

Multi-channel scalp EEG data was collected from both infant groups during day sleep. The requirement for the recording session was that each subject had to undergo two vigilance states: active sleep (AS) and quiet sleep (QS). EEG registration was performed using Waveguard caps with 19-28 sintered Ag/AgCl electrodes (ANT-Neuro, Berlin, Germany) located according to International 10-20 standard layout. Signals from both groups were recorded mostly with the NicOne EEG amplifier (Cardinal Healthcare, Ohio/Natus, Pleasanton, USA), but few EP subjects were recorded with the Cognitrace amplifier (ANT B.V., Enschede, The Netherlands). EEG recordings for both groups were performed at term-equivalent age of 41.4 ± 1.4 weeks CA (mean ± SD). The original sampling frequency was 256 Hz or 500 Hz, but all data was resampled to 250 Hz when exporting them into European Data Format (EDF).

### Clinical assessments

Newborn neurological assessments (Hammersmith Neonatal Neurological Examination (HNNE), (Dubowitz et al., 1999), were conducted on the EP and HC cohorts at term-equivalent age. The test comprises six separate domains of neurodevelopment: reflexes, movements, posture tonus, tone patterns, abnormal signs, orientation and behaviour. To render them suitable for studying associations with PPC networks, we used dimensionality reduction as described earlier (Tokariev et al., 2019b). In brief, we created combination scores, C1 and C2, using principal component analysis (PCA) with Varimax Rotation from three individual tests (visual alertness, head raising in prone, and increased neck extensor tone). In the post hoc assessment, the resulting C1 was shown to correlate primarily with later motor performance, whereas C2 was found to be associated with later cognitive and social performance (Tokariev et al., 2019b).

Long-term neurocognitive follow-up assessment was performed only on the EP infants at two years of age, using Bayley Scales of Infant and Toddler Development (Bayley, 2006) and the Griffiths Mental Developmental Scales (Huntley, 1996). The neuropsychological follow-up data was not available for the HC infants. These neurocognitive assessment tests were chosen, because of their established, widespread clinical use plus a broad impact on lifelong neurocognitive performance and quality of life, hence supporting the translational potential of findings (Hernandez, 2018; Rogers & Hintz, 2016). While some other outcomes such as gross motor development or hearing may be sometimes affected in EP infants, prior studies have suggested that these are most likely modified by a host of individual and treatment interventions (Kilbride et al., 2018).

### EEG review and pre-processing

Vigilance state assessment was performed through a combination of electrophysiological and behavioral measures, the latter observed using polygraphic channels (chin electromyogram, electrocardiogram, electrooculogram, and respiratory sensors). EEG traces during AS exhibit continuous fluctuations, with an irregular respiration and occasional eye movements. Conversely, EEG during QS is characteristically discontinuous, with a regular respiration (André et al., 2010). We then selected 5-min-long artifact-free EEG epochs from representative periods of AS and QS. To avoid transition periods between vigilance states, representative sleep epochs were selected from within well-established patterns of corresponding behavior and brain activity. Epochs that did not meet quality and length requirements were excluded from the data pool. As a result, we obtained four final groups: EP-AS (N = 46), HC-AS (N = 53), EP-QS (N = 42), and HC-QS (N = 66). For each subject we selected the same N = 19 channels (Fp1, Fp2, F7, F3, Fz, F4, F8, T7, C3, Cz, C4, T8, P7, P3, Pz, P4, P8, O1, O2) to enable group-level analysis. All EEG signals were first pre-filtered within the 0.15–45 Hz frequency range using a combination of high- and low-pass Butterworth filters of the 7th order. All filtering in this work were implemented offline and in forward-backward directions to compensate for phase delays introduced by infinite impulse response filters. Next, the EEG data were downsampled to a new sampling frequency, Fs = 100 Hz, and converted into average montage. Following our previous work (Tokariev et al., 2019a), we filtered the pre-processed EEG into 21 frequency bands of interest covering the range 0.4–22 Hz. Bandpass filtering was also implemented with pairs of low- and high-pass filters. The first central frequency (Fc) was set to 0.5 Hz and subsequent frequencies were computed as Fc(i) = 1.2×Fc(i-1), where i is the number of the frequency band. Cut-off frequencies for each band were taken as 0.85×Fc and 1.15×Fc correspondingly. This approach leads to 50% overlapping frequency bands of semi-equal width in the logarithmic scale.

### Computation of cortical signals

Band-pass filtered EEG were further source reconstructed to allow better spatial separation of cortical activities using a realistic infant head model (Tokariev et al., 2019b) and dynamic statistical parametric mapping (Dale et al., 2000). As the source space we used normal to cortical surface (at term age) dipoles of fixed orientation (N = 8014). The scalp and inner/outer skull shells were segmented from magnetic resonance imaging (MRI) data from healthy fullterm infant. Following previous studies (Despotovic et al., 2013; Odabaee et al., 2014; Tokariev et al., 2016a), tissue conductivities were set to: 0.43 S/m for scalp, 1.79 S/m for intracranial volume, and 0.2 S/m for skull. Finally, cortical sources were clustered into N = 58 parcels according to the scheme optimized for infant EEG. Cortical signals representing neural activity of each parcel were computed as the weighted mean of source signals belonging to the host parcels (Tokariev et al., 2019b).

### Computation of functional connectivity

To estimate functional connectivity, we computed phase-phase correlations (PPC) between all pairs of parcels using the debiased weighted phase-lag index (dwPLI), (Vinck et al., 2011). We opted to use this metric because of its robustness to artificial interactions caused by volume conduction (Palva et al., 2018; Palva & Palva, 2012). Connectivity was estimated using whole 5-min-long epochs, at 21 frequency bands and for both vigilance states (AS and QS). This led to a set of 58×58 PPC matrices for each subject. Next, we corrected these matrices by multiplication with a binary ‘fidelity’ mask, which was computed for the particular electrode layout. This mask was generated using extensive simulations based on the head model, and it removes ‘noisy’ connections from a connectivity matrix (Tokariev et al., 2019b). This procedure aims to improve the reliability of cortical-level network estimation from a suboptimal number of recording electrodes which are usually used in clinical recordings (Tokariev et al., 2016a). Note, that fidelity mask removes the same edges from all empirical connectivity matrices.

### Network analysis

To test network differences between EP and HC, we applied the Wilcoxon rank sum test (two one-tailed tests, α = 0.01) in an edge-by-edge manner with defined directions (EP > HC and EP < HC). This was done for each frequency band and for each sleep state separately. As a result of such edgewise scanning, we obtained matrices of *p*-values corresponding to the network connections and computed the ratio (*K*) of edges with significant group difference to the full network. To estimate the potential number of false discoveries in the frequency-specific group contrasts, we employed the Storey-Tibshirani adaptive FDR method using *q* = 0.01 with respect to the size of the whole network *(i.e.,* up to N = 1128 × 0.01 = 11 connections in each network were classified as potential false discoveries), (Puoliväli et al., 2020; Storey & Tibshirani, 2003). Effect size for the significantly different patterns was computed as a module of the mean of rank-biserial correlation values for corresponding connections. The influence of age differences on network strength was investigated by correlating the global mean connectivity strength with age for each group per frequency and sleep state (Spearman correlation, two-tailed test, α-level 0.05). The *p*-values of both sleep states were pooled together separately for each group and controlled for multiple comparisons by the Benjamini-Hochberg procedure (Benjamini & Hochberg, 1995).

Parallel to the primary analysis, a cross-check for statistical group comparison was performed using network-based statistics (NBS) (Zalesky et al., 2010) separately for the 21 frequency bands and two sleep states using two one-tailed tests (EP > HC and EP < HC). NBS is a multiple comparisons method designed specifically for network analysis. It assumes that connections reflecting true effects are interconnected into networks encompassing more than a single connection. The connected components are defined in a topological space, in contrary to other cluster-based methods, which use a physical space (Genovese et al., 2002; Zalesky et al., 2010). The initial threshold for the t-statistic was set to 2.5, followed by the post hoc permutation test to correct the family-wise error rate (5000 permutations, α = 0.05).

### Clinical correlation

The connectivity strength of each edge across the infant group was correlated with the corresponding neurological assessments at term-equivalent age and with the corresponding neurocognitive performance scores at two years of age (Spearman, two-tailed test with α-level 0.05) with conceptional age as a covariate. We computed the fraction of edges showing significant clinical correlation (*K*) for each frequency band and sleep state. Only the EP cohort had performance scores tested at 2 years of age. Some subjects had missing clinical scores, rendering the number of subjects used for each correlation to: C1: EP-AS (N = 39), EP-QS (N = 36), HC-AS (N = 30), HC-QS (N = 51); C2: EP-AS (N = 39), EP-QS (N = 36), HC-AS (N = 40), HC-QS (N = 51); Griffiths Visual: EP-AS (N = 35), EP-QS (N = 32); Griffiths Motor: EP- AS (N = 39), EP-QS (N = 36); Bayley Cognitive: EP-AS (N = 32), EP-QS (N = 30); Bayley Language comprehension: EP-AS (N = 30), EP-QS (N = 28). Multiple comparisons correction was implemented with the Storey-Tibshirani adaptive FDR with *q* = 0.05 *(i.e.,* 2.5 % of the positive and negative correlations separately were classified as potential false discoveries). The Spearman *ρ*-value was used for estimating effect size: it was computed for all significant edges and averaged across the full network.

### Analysis software

Source reconstruction was conducted using the Brainstorm (Tadel et al., 2011), (https://neuroimage.usc.edu/brainstorm/Introduction), and the openMEEG (Gramfort et al., 2010), (https://openmeeg.github.io/), software packages. Analyses were performed with Matlab R2020a (MathWorks, Naticks, MA, USA) and NBS Connectome (Zalesky et al., 2010), (https://www.nitrc.org/projects/nbs/), and the visualization of brain networks was carried out with BrainNet Viewer (Xia et al., 2013), (https://www.nitrc.org/projects/bnv/).

The Matlab script implementing the network analyses of group differences and clinical correlation can be found at https://github.com/pauliina-yrjola/Preterm-Phase.

## Acknowledgements

This work was supported by the Finnish Pediatric Foundation, the Finnish Academy (313242, 288220, 321235), Juselius Foundation, Aivosäätiö, Neuroscience Center at University of Helsinki, as well as Helsinki University Central Hospital

## Supplementary Figures

**Figure 2—figure supplement 1.**
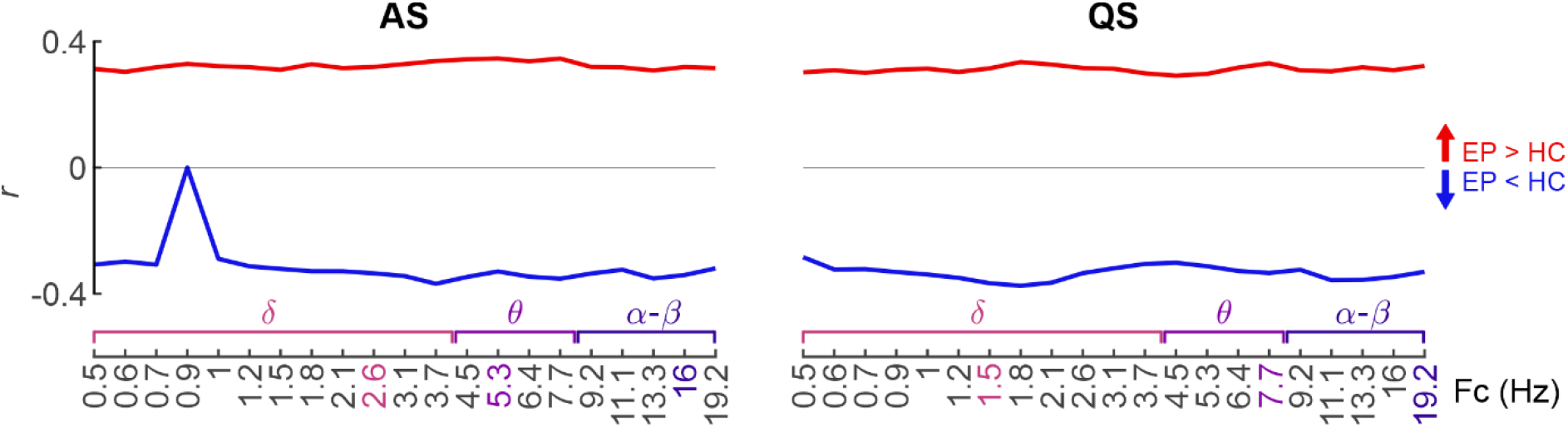
Effect size of the statistical group differences on cortical PPC networks. The effect size, computed by the mean rank-biserial correlation (*r*) of the significant networks shown in Figure 1 (two one-tailed Wilcoxon rank sum tests, α = 0.01) during active sleep (AS, left) and quiet sleep (QS, right), as a function of frequency. Networks with increased connectivity strength in EP (EP > HC) are shown in red, and networks with decreased connectivity in EP (EP < HC) are displayed in blue.

**Figure 2—figure supplement 2.**
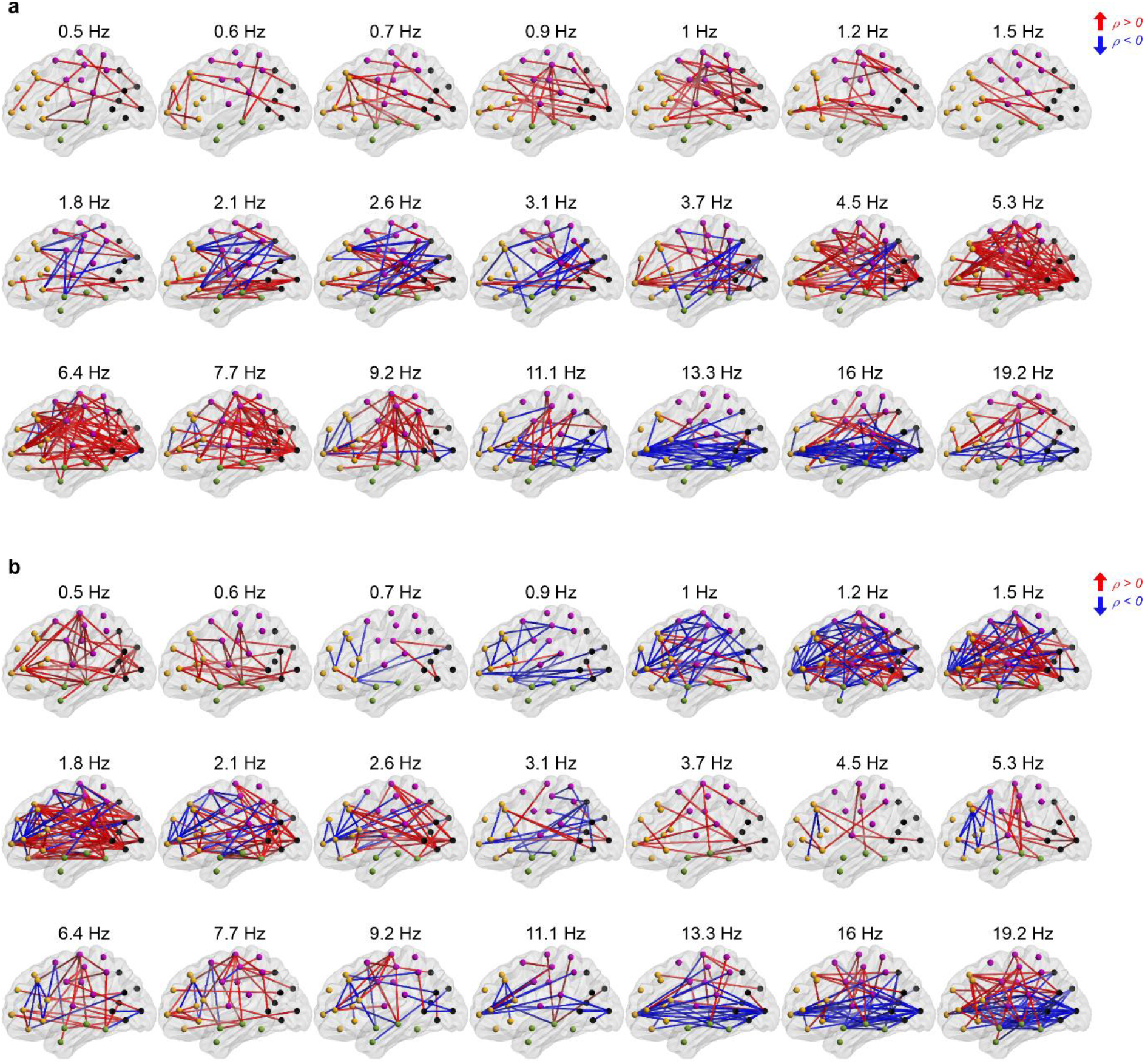
Effects of prematurity on cortical PPC networks modulate over frequency. Spatial visualizations of the group difference networks obtained from network density measurements (Figure 1, two one-tailed Wilcoxon rank sum tests, α = 0.01) over all frequency bands in AS (a) and QS (b). Only the edges which passed FDR correction (*q* = 0.01) are shown. Red networks display connections of increased connectivity in EP (EP > HC) and blue networks reduced connectivity in EP (EP < HC).

**Figure 2—figure supplement 3.**
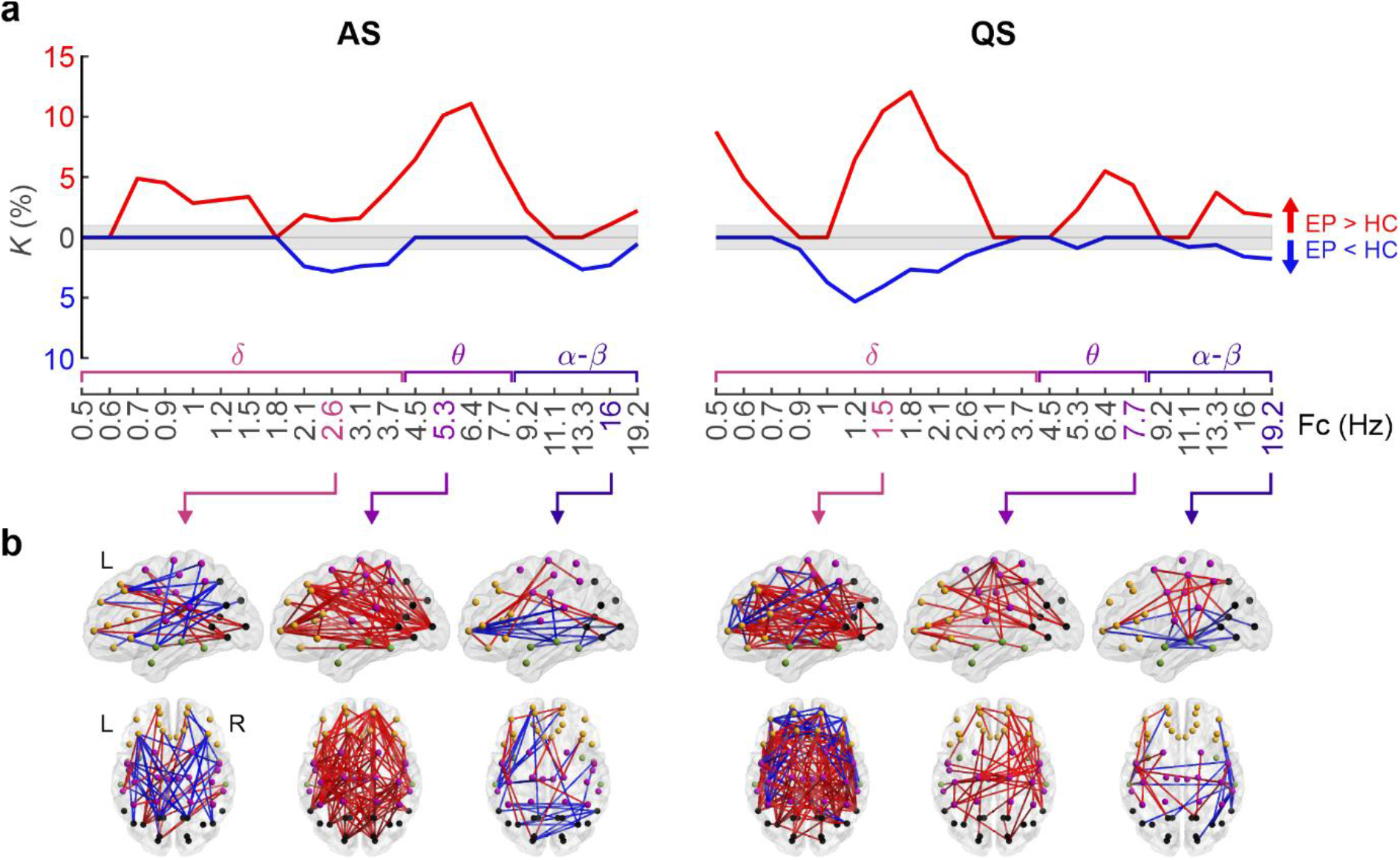
Effects of prematurity on cortical PPC networks replicated with an independent additional analysis. **(a)** Network density *(K)* of PPC group difference networks computed with NBS Connectome (two one-tailed tests, threshold 2.5, significance level α = 0.05, 5000 permutations) is shown for AS (left) and QS (right) as a function of frequency bands, denoted by their central frequencies. The directions of difference (EP > HC or EP < HC) are indicated in red and blue, respectively. **(b)** 3-dimensional visualizations present the spatial distribution of group difference networks at the frequencies selected from Figure 1. The direction of difference is indicated as in (a).

**Figure 3—figure supplement 1.**
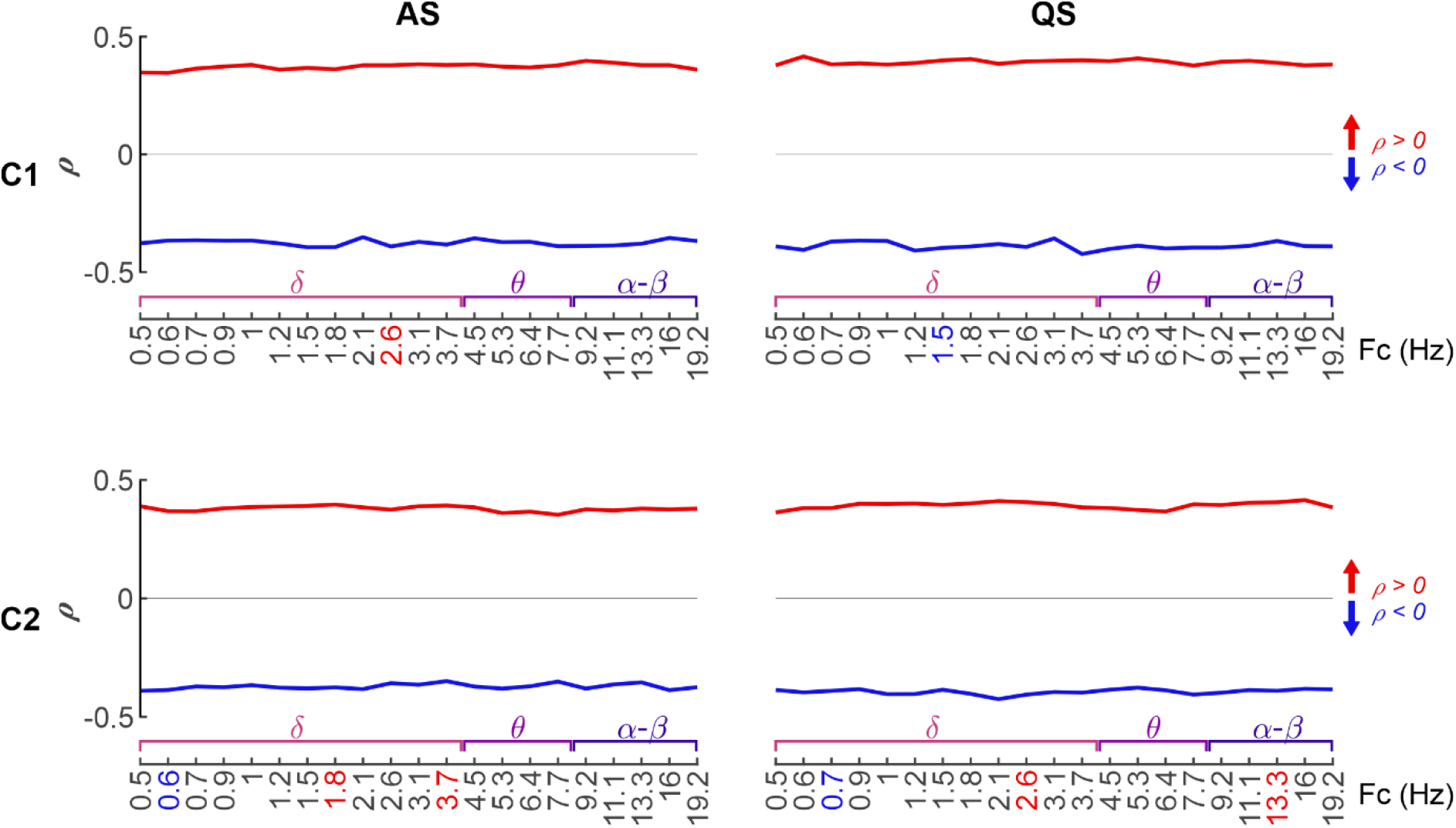
Effect size of PPC networks depicting clinical correlation. The effect size, acquired from the mean Spearman (two-tailed test, with conceptional age as a covariate, and α = 0.05) ρ-value of the positive (ρ ≥ 0, red) and negative (ρ < 0, blue) networks, as a function of frequency band. The results are presented for active sleep (AS, left) and quiet sleep (QS, blue), as well as for the neurological outcome scores C1 (above) and C2 (below).

**Figure 3—figure supplement 2.**
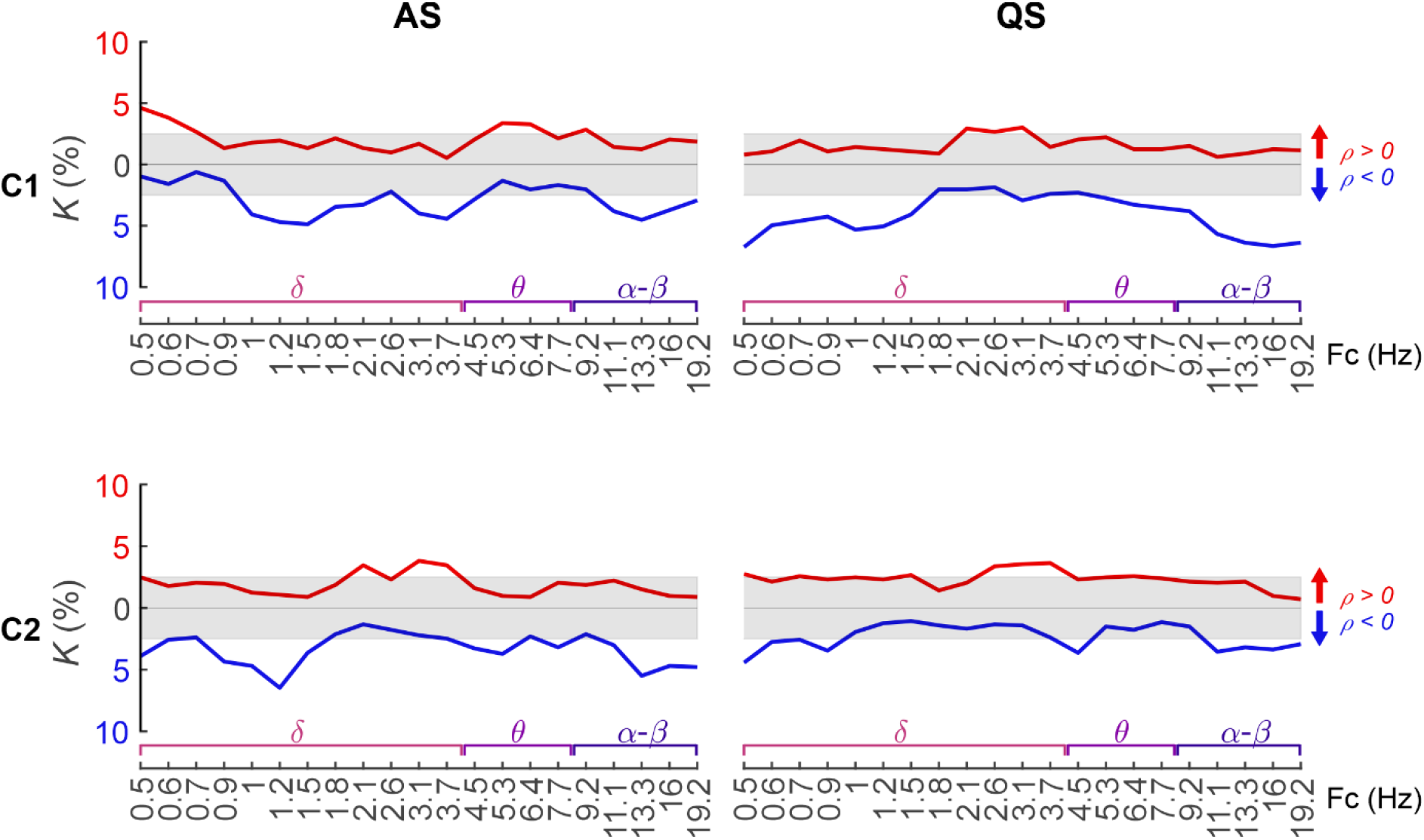
Absence of correlation between cortical PPC strengths and early neurological performance in healthy controls. Network density *(K)* of PPC correlation related to the neurological assessment scores C1 and C2 as a function of frequency band in the HC cohort (Spearman, two-tailed test, with conceptional age as a covariate, and α = 0.05). The FDR (*q* = 0.05) boundaries are depicted as a grey shaded area. Colour coding represents the polarity of the correlation (red: ρ ≥ 0, blue ρ < 0). Analysis of the HC group shows an absence of wider network patterns that would correlate positively to either of the neurological scores. There are some networks at low frequencies with negative correlation to C1 and C2 during AS (peak at Fc = 1.2-1.5 Hz) and some networks at low and high frequencies with negative correlation to C1 during QS (peaks at Fc = 0.5 Hz and Fc = 16 Hz).

**Figure 3—figure supplement 3.**
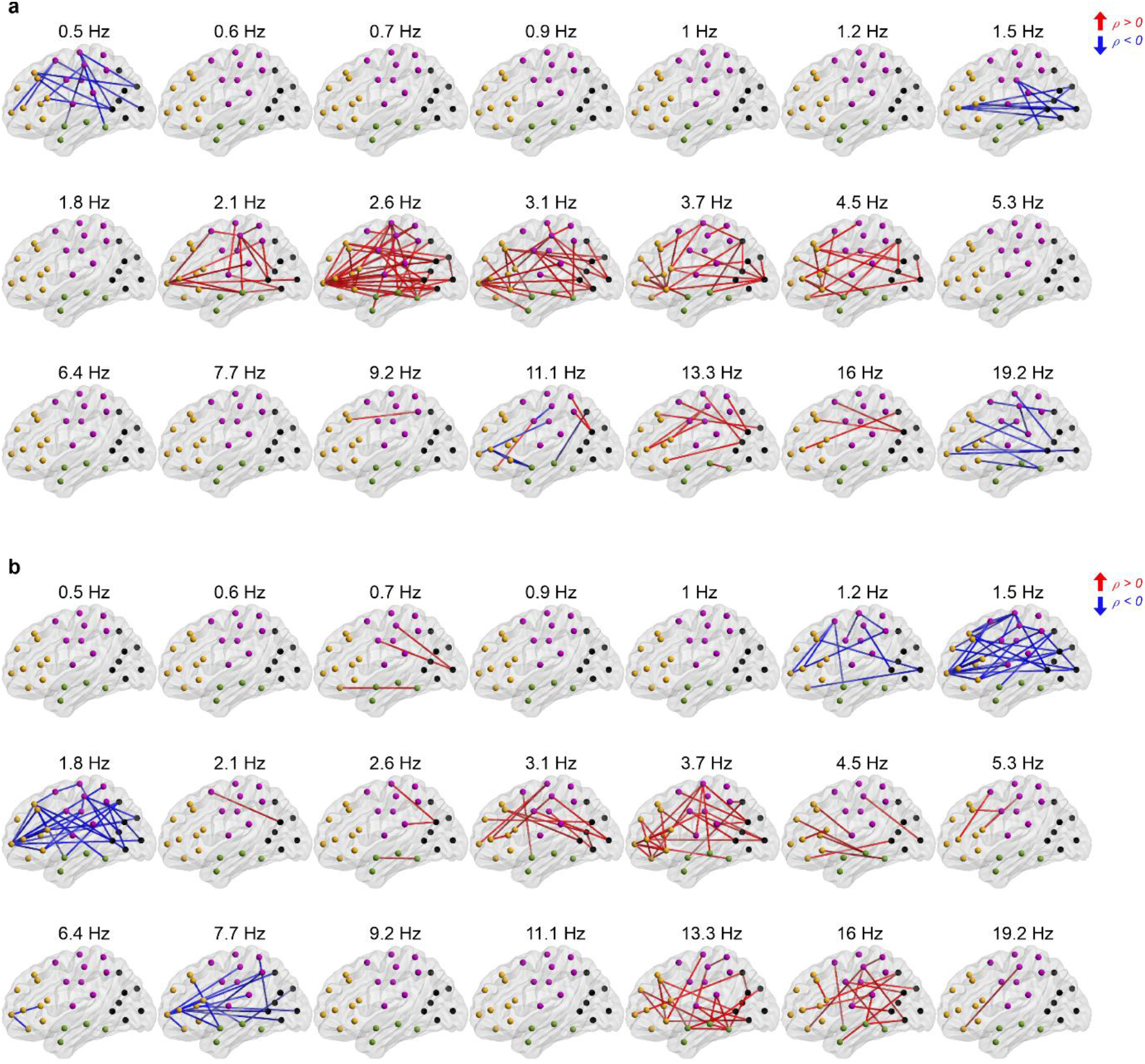
The frequency-specific PPC correlation networks to C1 neurological outcomes. 3-dimensional visualizations depicting the correlation of connection strength to C1 scores (Spearman, two-tailed test, with conceptional age as a covariate, and α = 0.05) on all investigated frequency bands in AS (a) and QS (b) in EP infants at term-equivalent age. The edges displayed in the figure passed FDR correction (*q* = 0.05). Red networks indicate positive correlation (Spearman ρ ≥ 0), whereas blue connections express negative correlation (Spearman ρ < 0).

**Figure 3—figure supplement 4.**
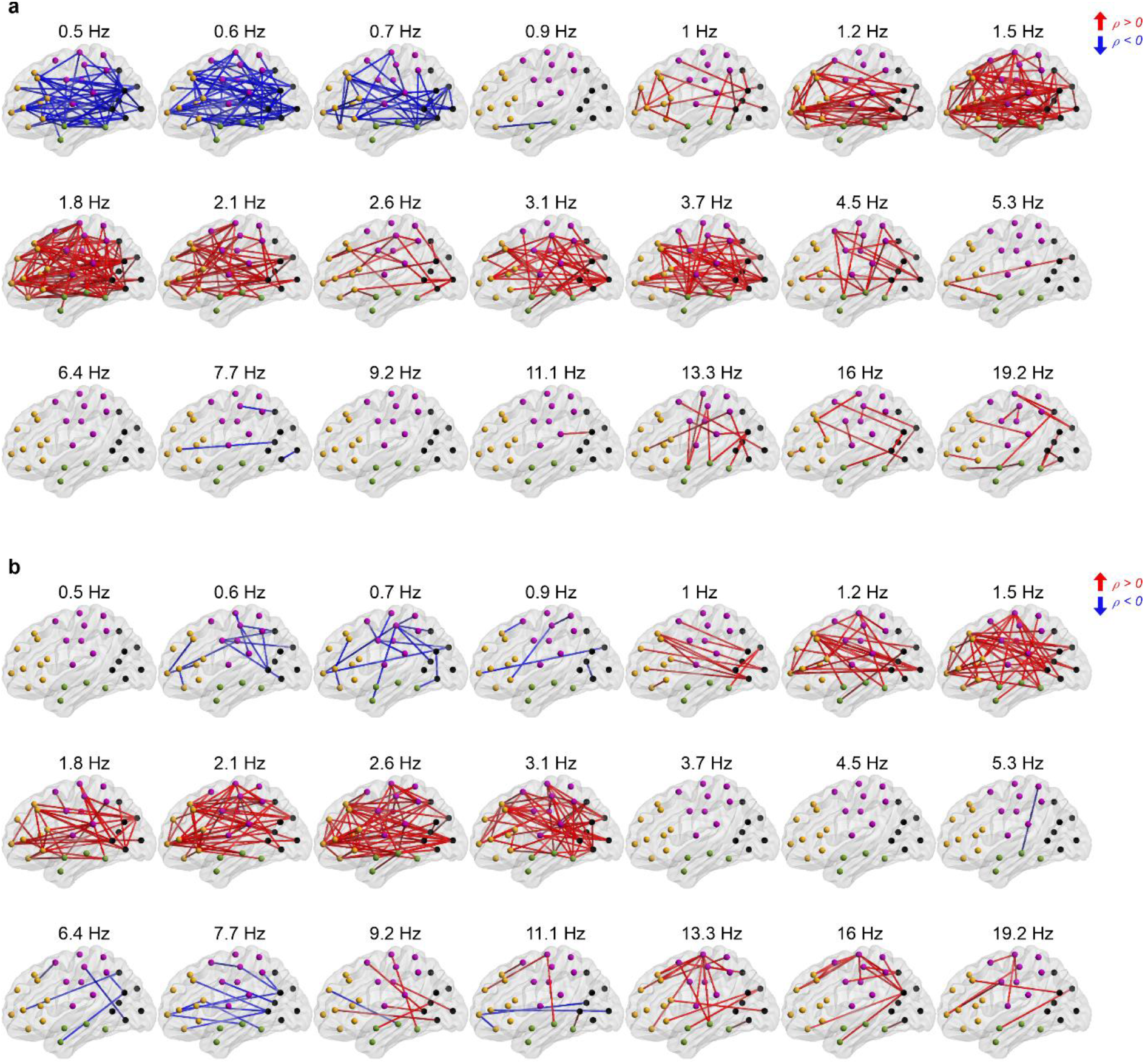
The frequency-specific PPC correlation networks to C2 neurological outcomes. 3-dimensional visualizations depicting the correlation of connection strength to C2 scores (Spearman, two-tailed test, with conceptional age as a covariate, and α = 0.05) on all investigated frequency bands in AS (a) and QS (b) in EP infants at term-equivalent age. The edges displayed in the figure passed FDR correction (*q* = 0.05). Red networks indicate positive correlation (Spearman ρ ≥ 0), whereas blue connections express negative correlation (Spearman ρ < 0).

**Figure 4—figure supplement 1.**
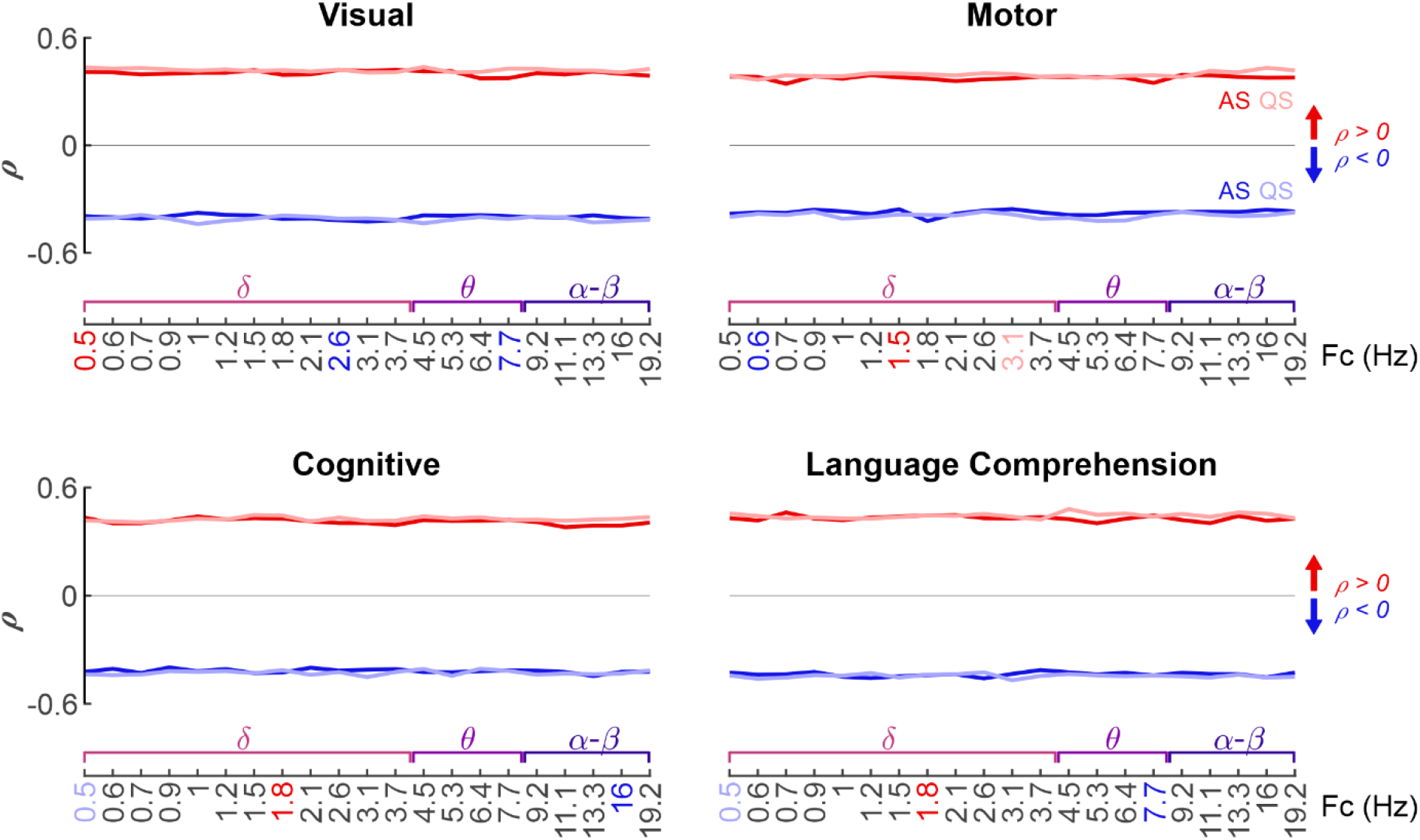
Effect size of PPC networks depicting long-term neurocognitive correlation. The effect size was computed as the mean Spearman (twotailed test, with conceptional age as a covariate, and α = 0.05) ρ-value for the significant networks of each frequency band. The colors show the sign of the correlation (ρ ≥ 0: red and ρ < 0: blue). The effect size values are presented separately for active sleep (AS, dark hues) and quiet sleep (QS, light hues) as well as for the neurocognitive scores Griffiths visual (upper left), Griffiths motor (upper right), Bayley cognitive (lower left), and Bayley language comprehension (lower right).

**Figure 4—figure supplement 2.**
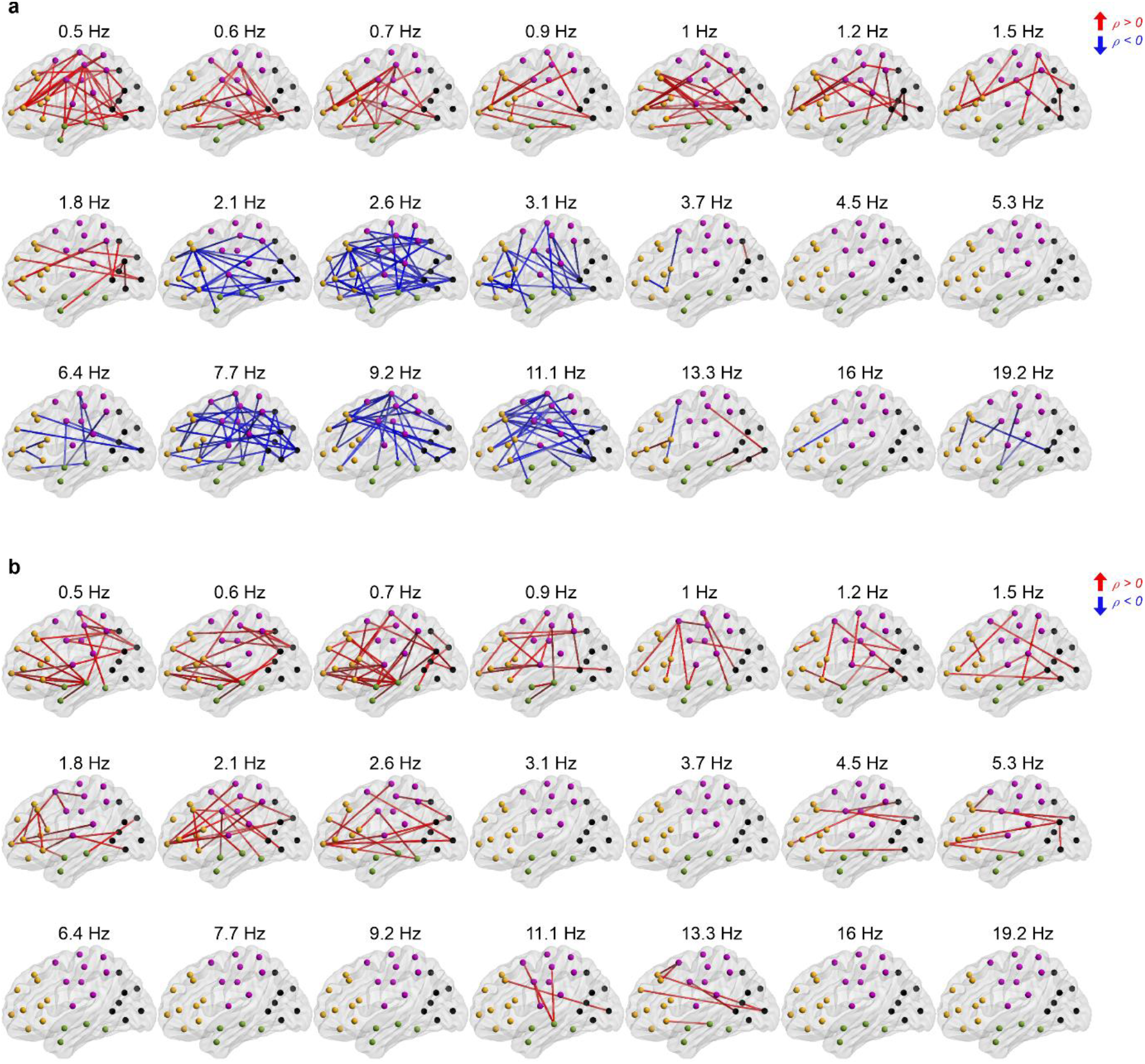
The frequency-selective PPC fingerprint networks reflecting visual performance at 2 years of age. Spatial visualizations of PPC edge strength correlation to Griffiths visual scores (Spearman, two-tailed test, with conceptional age as a covariate, α = 0.05) over all examined frequency bands in AS (a) and QS (b) in the EP cohort at 2 years of age. The presented connections survived multiple comparisons correction with FDR (*q* = 0.05). Colour coding represents the sign of correlation (red: ρ ≥ 0, blue ρ < 0).

**Figure 4—figure supplement 3.**
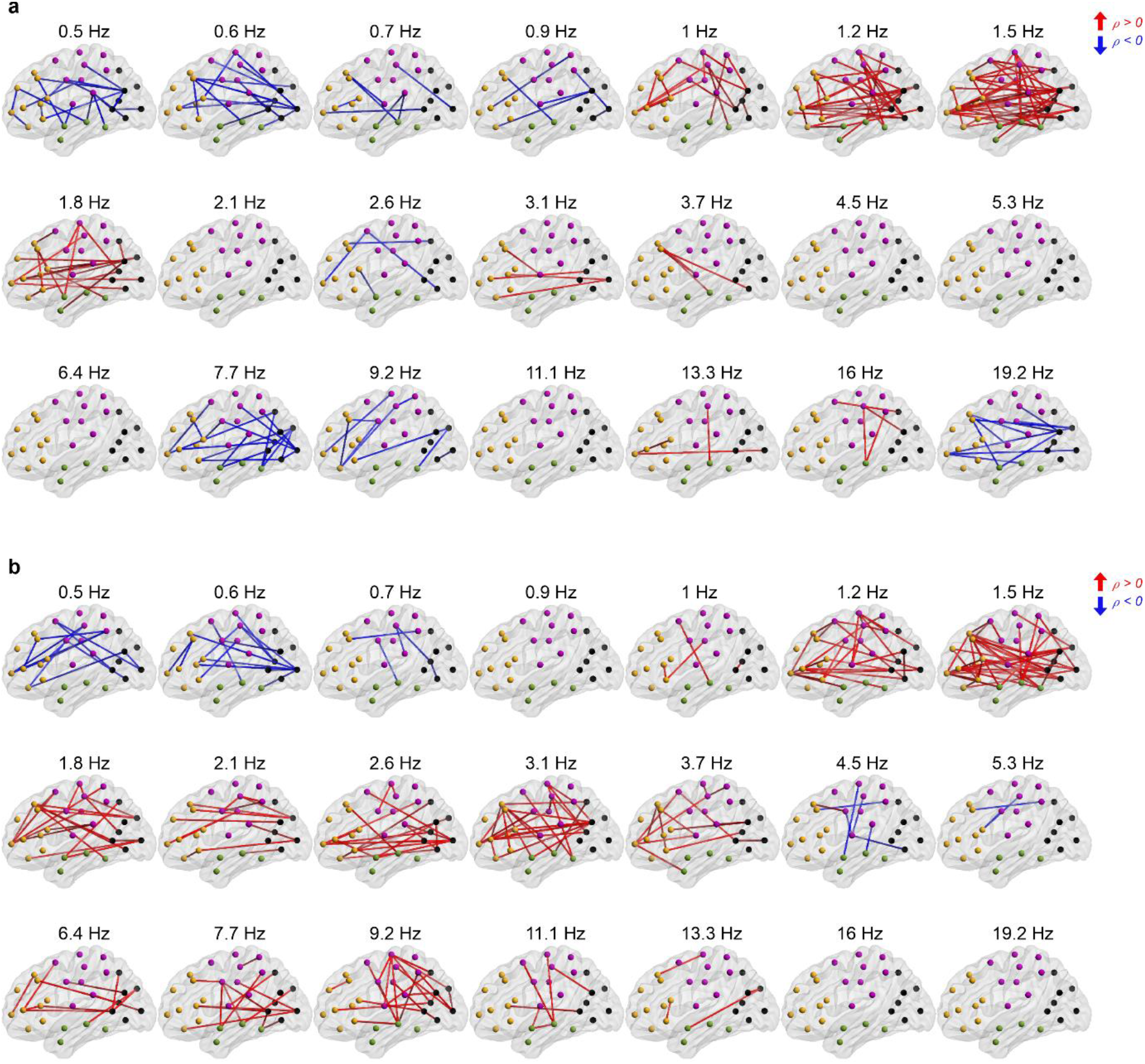
The frequency-selective PPC fingerprint networks reflecting motor performance at 2 years of age. Spatial visualizations of PPC edge strength correlation to Griffiths motor scores (Spearman, two-tailed test, with conceptional age as a covariate, α = 0.05) over all examined frequency bands in AS (a) and QS (b) in the EP cohort at 2 years of age. The presented connections survived multiple comparisons correction with FDR (*q* = 0.05). Colour coding represents the sign of correlation (red: ρ ≥ 0, blue ρ < 0).

**Figure 4—figure supplement 4.**
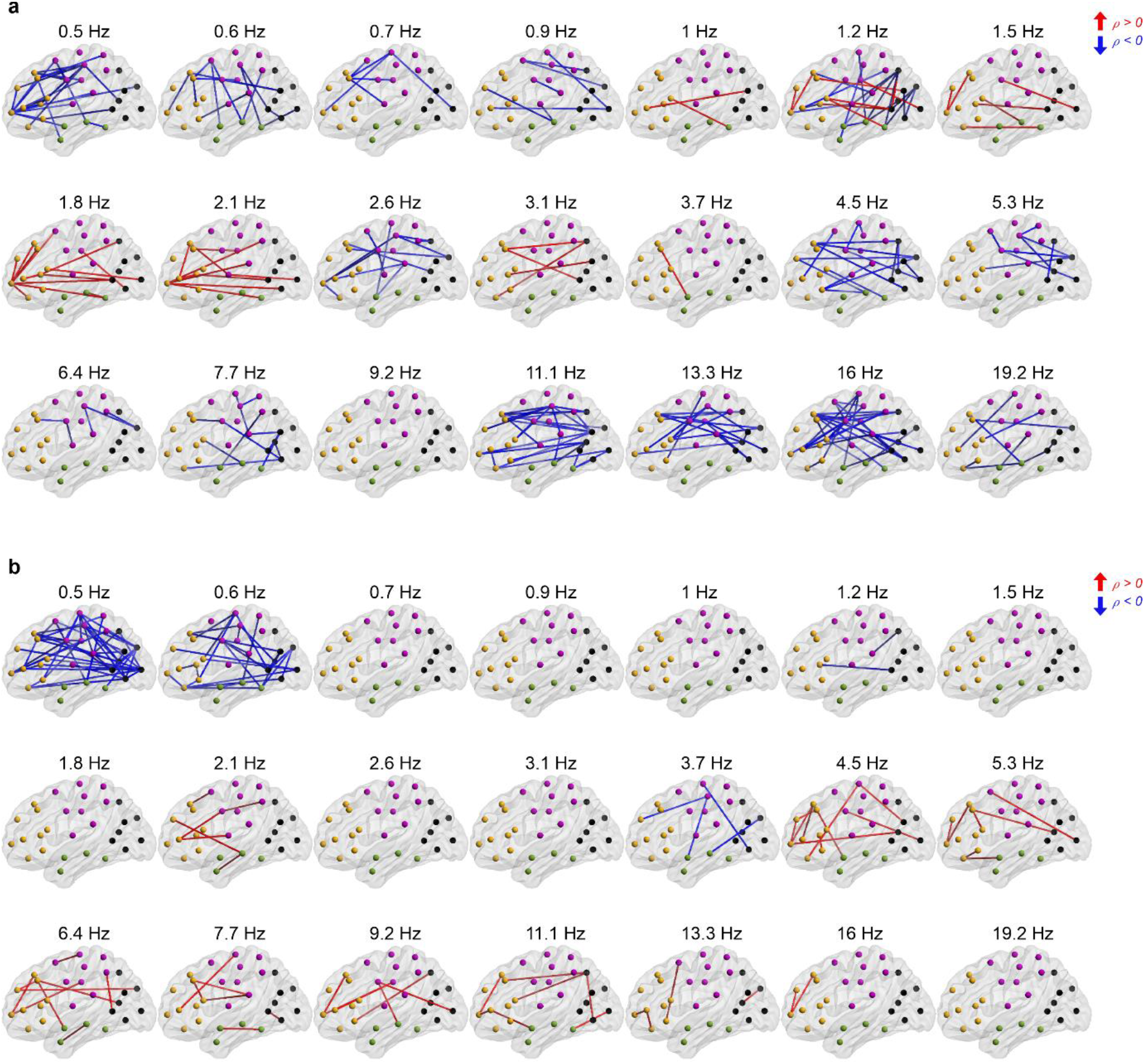
The frequency-selective PPC fingerprint networks reflecting cognitive performance at 2 years of age. Spatial visualizations of PPC edge strength correlation to Bayley cognitive scores (Spearman, two-tailed test, with conceptional age as a covariate, and 0.05) over all examined frequency bands in AS (a) and QS (b) in the EP cohort at 2 years of age. The presented connections survived multiple comparisons correction with FDR (*q* = 0.05). Colour coding represents the sign of correlation (red: ρ ≥ 0, blue ρ < 0).

**Figure 4—figure supplement 5.**
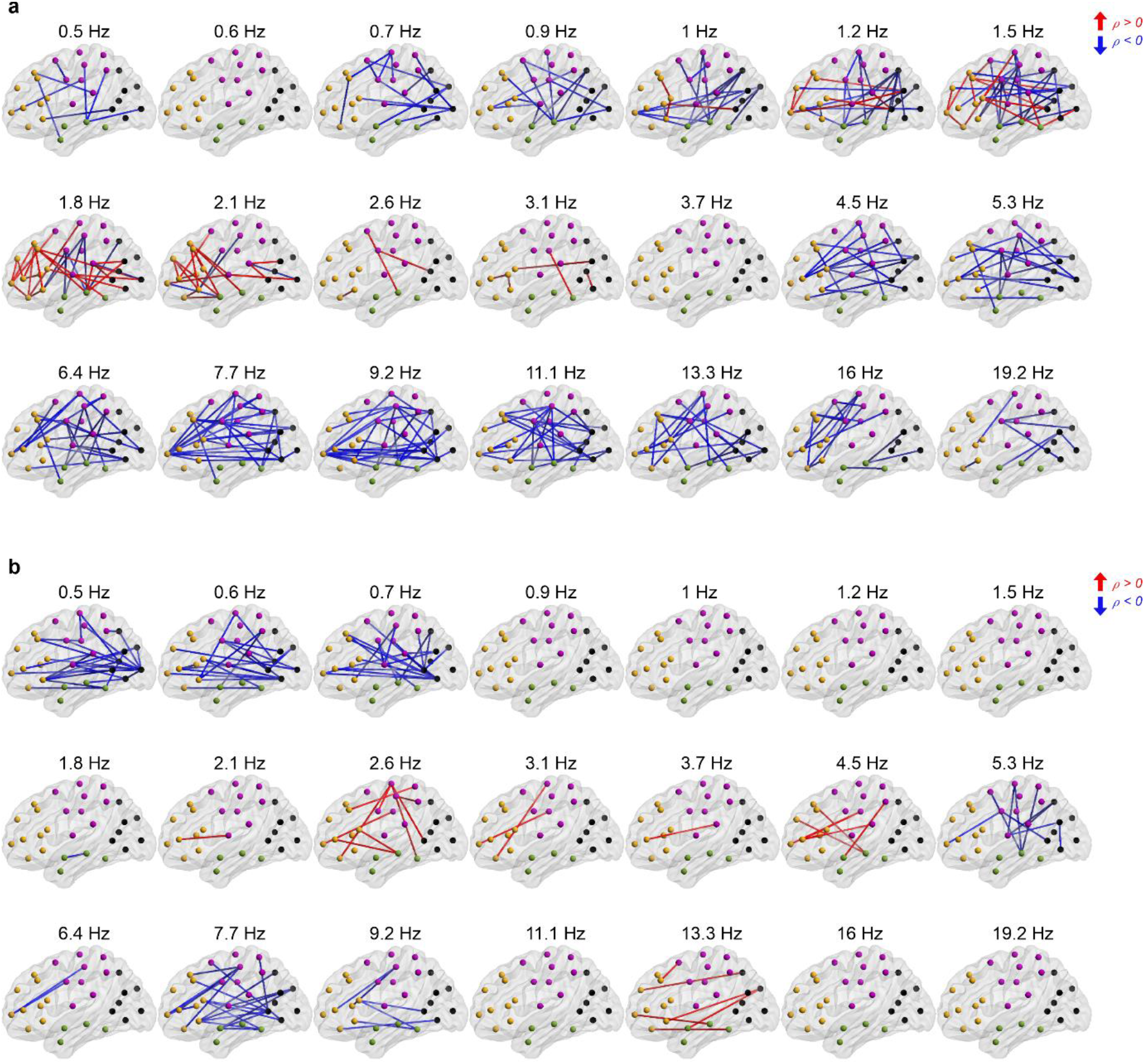
The frequency-selective PPC fingerprint networks reflecting language comprehension at 2 years of age. Spatial visualizations of PPC edge strength correlation to Bayley language comprehension scores (Spearman, two-tailed test, with conceptional age as a covariate, α = 0.05) over all examined frequency bands in AS (a) and QS (b) in the EP cohort at 2 years of age. The presented connections survived multiple comparisons correction with FDR *(q* = 0.05). Colour coding represents the sign of correlation (red: ρ ≥ 0, blue ρ < 0).

